# Host range and ARG dissemination are shaped by distinct survival strategies of conjugative plasmids

**DOI:** 10.1101/2025.05.22.655438

**Authors:** Ryuichi Ono, Naoki Konno, Yuki Nishimura, Chikara Furusawa

**Affiliations:** Innovative Genomics Institute, University of California, Berkeley, Berkeley, CA 94720, USA; Department of Comparative Biochemistry, University of California, Berkeley, Berkeley, CA 94720, USA; Department of Integrated Biosciences, Graduate School of Frontier Sciences, the University of Tokyo, Kashiwa, Chiba, 277-0882, Japan; Department of Biological Sciences, Graduate School of Science, the University of Tokyo. Bunkyo-ku, Tokyo, 113-0032, Japan; Universal Biology Institute, The University of Tokyo, 7-3-1 Hongo, Bunkyo-ku, Tokyo 113-0033, Japan; Center for Biosystems Dynamics Research, RIKEN, 6-7-1 Minatojima-minamimachi, Chuo-ku, Kobe, 650-0047, Japan

**Author notes:** To whom correspondence should be addressed. Correspondence may also be addressed to Naoki Konno. (R.O.), (N.K.); Fax: Not Available; (R.O.), (N.K.).

## Abstract

Horizontal gene transfer is a major driver of bacterial evolution and the global dissemination of antibiotic resistance genes (ARGs). Conjugative plasmids play a crucial role in ARG spread across hosts within their host range, yet the genetic and functional determinants shaping plasmid host range remain poorly understood. Here, we systematically analyzed the gene content of conjugative/mobilizable plasmids from public databases and found that two distinct survival strategies were enriched in different host-range groups: a “stealth” strategy, which minimizes host fitness costs by employing a global regulator *h-ns*, was particularly enriched in broad-host-range plasmids, whereas an “aggressive” strategy, which protects plasmids by actively inhibiting host response including SOS response by encoding the SOS inhibitor *psiB*, was more common in narrow-host-range plasmids. Plasmids employing either strategy constituted the majority of all conjugative plasmids analyzed, and accumulated significantly more ARGs than plasmids with neither strategy. Our data further suggested that stealth plasmids facilitate the acquisition of emerging ARGs, while aggressive plasmids amplify the copy number of established ARGs. This “stealth-first” model successfully recapitulated historical ARG dissemination patterns. These findings provide critical insights into the relationship between plasmid survival strategies and host range, advancing our understanding of the global patterns underlying plasmid-mediated ARG transmission.

**GRAPHICAL ABSTRACT:** 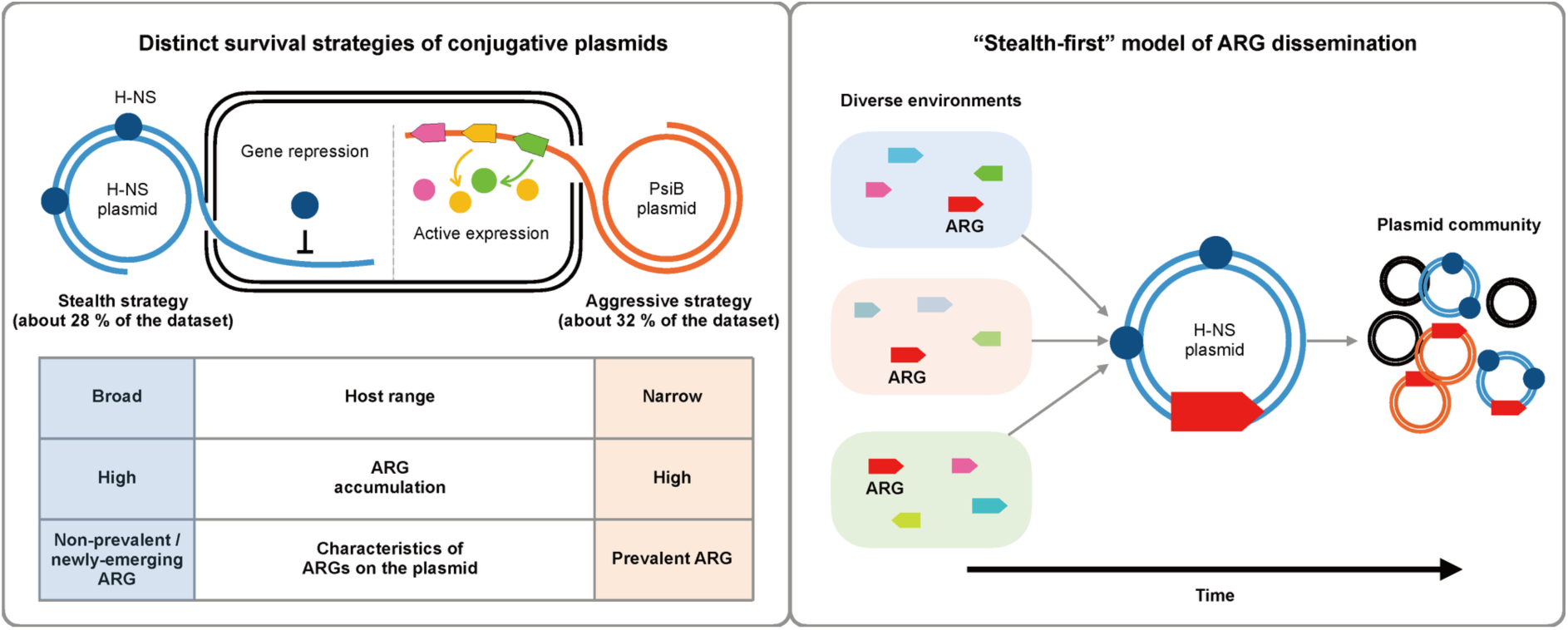

## INTRODUCTION

Horizontal gene transfer (HGT) plays a pivotal role in the remarkable adaptability and rapid evolution of prokaryotes (1, 2). A significant proportion of HGT events are mediated by mobile genetic elements (MGEs), such as plasmids. HGT is a complex phenomenon driven by a trade-off between the selfish behavior of MGEs, which exploit host resources to increase their own copy number, and the potential benefits to the host from acquiring new genes (e.g., antibiotic resistance genes). Previous studies have demonstrated that HGT is generally infrequent between evolutionarily distant host species or across disparate environments (3–5). Additionally, the occurrence of HGT is often constrained by factors such as the presence of specific defense systems within hosts or abnormal protein-protein interactions (PPIs) between transferred and host proteins (6–8). However, little is known about what genetic traits or adaptive mechanisms make certain MGEs particularly prone to facilitating HGT in natural environments. Understanding these mechanisms is not only essential for leveraging MGEs as tools for DNA delivery (9, 10) but also critically important for elucidating the emergence and spread of antibiotic resistance, an urgent global challenge.

According to estimates by the WHO, antibiotic-resistant bacteria were responsible for approximately 1.27 million deaths in 2019 (11). Without appropriate measures, this number is predicted to swell to 10 million deaths annually by 2050 (12). Furthermore, the World Bank has stated that the potential direct and indirect economic damage would be significant, with annual costs potentially rivaling those of the global financial crisis that began in 2008 (13). The dissemination and acquisition of antibiotic resistance genes (ARGs) are heavily influenced by HGT mediated by MGEs (14). Among these, conjugative plasmids— extrachromosomal DNA elements capable of transmitting between cells via conjugation— and mobilizable plasmids, which lack some components of the conjugation machinery but can be transferred using machinery provided by other MGEs, have been critical focal points of research. This is due to their significant role in the accumulation of ARGs (15, 16) and the ability to produce novel ARGs through mutagenesis triggered by conjugation (17, 18). Another critical feature that underscores the importance of conjugative plasmids in ARG dissemination is their potential to broaden their host ranges. Some conjugative plasmids are capable of being transferred across phylogenetically distant species, even across phyla (19, 20) or domains (21, 22). Such broad host range often exceeds that of bacteriophages, which typically rely on highly specific (e.g. species- or strain-specific) interactions with host receptors for successful infection (23). ARGs encoded on these broad-host-range MGEs have the potential to spread not only across different bacterial species but also across diverse biological and ecological contexts, including human and animal hosts as well as environmental niches. As a result, they represent one of the most critical factors in the global dissemination of ARGs (24).

Against this backdrop, broad-host-range (BHR) plasmids remain a fascinating research focus to explore the dynamics of ARG dissemination. Recent advancements in computational methods have enabled more precise classification of plasmids and improved predictions of their host range (20). Concurrently, various studies have revealed correlations between host range and genomic features, such as the number of restriction sites (25), and the number and structure of replication origins (26), suggesting that plasmids with different host ranges may require distinct strategies for adaptations. However, considering that plasmids serve as natural vehicles for gene transfer, a critical but largely unresolved question is how plasmid-encoded gene repertoires correspond to host range. Furthermore, the survival strategies inferred from these gene repertoires and their implications for ARG dissemination remain largely unexplored.

In this study, we performed a comprehensive gene enrichment analysis on plasmids with previously annotated host ranges to systematically investigate how conjugative plasmids shape their host range and contribute to ARG dissemination. We identified two genes, *h-ns* and *psiB*, that were strongly associated with BHR and narrow-host-range (NHR) plasmids, respectively, and found to be almost mutually exclusive. Remarkably, plasmids carrying either gene represented the majority of conjugative/mobilizable plasmids in our dataset and exhibited a significantly greater tendency to accumulate ARGs than those lacking both. Further analyses revealed that H-NS plasmids—associated with a “stealth” strategy that minimizes host fitness costs—tend to acquire newly emerging ARGs, while PsiB plasmids— employing an “aggressive” strategy that suppresses host stress responses—preferentially disseminate ARGs already prevalent in microbial communities. A temporal analysis of 48 major ARGs uncovered a consistent, global trend in which novel ARGs are first acquired by stealth plasmids and subsequently spread via aggressive plasmids. This sequential dynamic supports a proposed “stealth-first” model of plasmid-mediated ARG dissemination. Together, these findings unveil distinct survival strategies that underpin plasmid host range and ARG dynamics, offering a new conceptual framework for understanding and predicting the evolutionary trajectories of antibiotic resistance.

## MATERIAL AND METHODS

### Datasets and gene annotation

A total of 943 sequences from Enterobacterales were used for the enrichment analysis. These sequences were derived from the 84th NCBI RefSeq database (27) and had been assigned MOB types and host range (Hrange) classifications in a prior study (20). Additionally, on October 5, 2024, the PLSDB database (version: 2021_06_23_v2) (28, 29) was downloaded and used for subsequent analyses. For both datasets, GenBank files were retrieved via the NCBI Entrez API using the accession numbers indicated in the metadata files. For each CDS from the GenBank files, gene annotation was performed by running HMMscan from HMMER (version: 3.3) (30) against PFAM r35 with an E-value cutoff of 10⁻² (-E 1e-2). Plasmids were categorized as H-NS plasmid, PsiB plasmid, Both plasmid or None plasmid based on the presence or absence of genes annotated as “Histone_H-NS” (PF00816.24) and “PsiB” (PF06290.14).

### Enrichment analysis

For the 943 sequences from Enterobacterales with MOB type and Hrange information, a previous study(20) assigned Hrange values based on the plasmid host range. ‘Single species’ plasmids, which have the narrowest host range (Hrange = ‘I’), are restricted to a single bacterial species. Plasmids capable of transferring to multiple species but limited to those within the same family were assigned Hrange = ‘II’. Similarly, plasmids with Hrange values of ‘III’, ‘IV’, ‘V’, and ‘VI’ were defined based on increasing host range breadth. The broadest category, Hrange = ‘VI’, includes plasmids that can transfer to bacteria spanning multiple phyla. Plasmids with an Hrange of ‘I’, ‘II’, or ‘III’ were classified as narrow-host-range (NHR) plasmids (701 sequences), while those with an Hrange of ‘IV’, ‘V’, or ‘VI’ were classified as broad-host-range (BHR) plasmids (242 sequences). To compare the frequency of each detected gene between NHR and BHR plasmids, Fisher’s exact test (fisher_exact from SciPy, version: 1.14.0) was conducted using HMMscan results. False Discovery Rate (FDR) for each hit was calculated by Benjamini-Hochberg procedure (false_discovery_control from SciPy, version: 1.14.0). To assess the statistical significance of the mutually exclusive distribution of *psiB* and *h-ns*, Chi-squared test (chi2_contingency from SciPy, version: 1.14.0) was conducted. Additionally, the Jaccard index was calculated for all pairs of genes encoded by more than 250 plasmids.

The goal of this analysis was to identify genes that characterize narrow or broad host ranges. However, enrichment analysis using all genes detected by HMMscan risks identifying genes specific to certain MOB types or even smaller subgroups, which may not reflect host range characteristics. To address this, the analysis was restricted to genes present in at least 10% of plasmids within each host range type and detected across a diverse set of MOB types. To measure the MOB-type distribution diversity of each gene, Simpson’s index was employed. The histograms of Simpson’s index for genes detected in BHR and NHR plasmids revealed three peaks near 0, 0.2, and 0.4 (Supplementary Fig. 2). The peak near 0 likely corresponds to genes specific to small plasmid subgroups within each host range, while the peaks near 0.2 and 0.4 represent genes more broadly distributed across plasmids within the same host range. To further investigate the distribution, we plotted the proportion of genes with Simpson’s index values exceeding a threshold n (0 ≤ n ≤ 1). A clear drop was observed near n=0.4 in both BHR and NHR plasmids (Supplementary Fig. 2). Based on this observation, a Simpson’s index threshold of 0.4 was applied to identify genes broadly distributed across plasmids for subsequent analyses.

### Filtering and Preliminary Characterization of Conjugative Plasmids from PLSDB

All plasmids in PLSDB (version: 2021_06_23_v2) were analyzed using MOB-typer (version: 3.1.9) (31). Plasmids classified as “conjugative” with sequence lengths of 500 kb or less were used for downstream analyses (hereafter referred to as PLSDB plasmids). Sequence length and GC content were obtained from metadata provided by PLSDB. IS content (%) was calculated for all PLSDB plasmids using ISEScan (version: 1.7.2.3) (32). Chi-squared test (chi2_contingency from SciPy, version: 1.14.0) was conducted to assess the statistical significance of the mutually exclusive distribution of PsiB and H-NS. To compare the distribution of GC content, length, IS content, ARG number per 100 kb, and integron content, Mann-Whitney U test (mannwhitneyu from SciPy, version: 1.14.0) was conducted. False Discovery Rate (FDR) was calculated by Benjamini-Hochberg procedure (false_discovery_control from SciPy, version: 1.14.0). Additionally, FastANI (version: 1.33) (33) and ANIclustermap (version: 1.4.0) (34) was used to compute the Average Nucleotide Identity (ANI) for all possible pairs in PLSDB plasmids. A dendrogram based on ANI was visualized using iTOL v7 (35).

### Annotating leading region

Using the results of MOB-typer, PLSDB plasmids containing single oriT and single relaxase were filtered for further analysis. Among these, plasmids in which the oriT and relaxase were located nearby (within 1, 000 bp) (36) were classified as having a putative leading region. Clinker (37) was used for the visualization of example-leading regions. To compare the distribution of plasmids with the leading region, Chi-squared test (chi2_contingency from SciPy, version: 1.14.0) was conducted. False Discovery Rate (FDR) was calculated by Benjamini-Hochberg procedure (false_discovery_control from SciPy, version: 1.14.0).

### ARG and integron annotation

AMRFinderPlus (version: 3.12.8, AMRFinder database version: 2024-07-22.1) (38) was used for PLSDB plasmids to detect ARGs. Only ARGs with the annotation of ‘Element type = AMR’ were included in downstream analyses. Integron regions within each plasmid were annotated using IntegronFinder 2.0 (39). Detected ARGs were classified into three categories based on their prevalence in the dataset. ARGs encoded by fewer than 10 plasmids were classified as ‘Rare’, those found in 10 to 99 plasmids as ‘Moderate’, and those present in 100 or more plasmids as ‘Prevalent’. For each ARG, if its start and end positions were entirely contained within an integron region, it was annotated as an ARG encoded within an integron. To compare the distribution of the Proportions of plasmid types carrying each ARG, Wilcoxon rank-sum test (wilcoxon from SciPy, version: 1.14.0) was conducted. False Discovery Rate (FDR) was calculated by Benjamini-Hochberg procedure (false_discovery_control from SciPy, version: 1.14.0). To compare the distribution of ‘Rare’ and ‘Moderate’ ARGs across different plasmid types, Chi-squared test (chi2_contingency from SciPy, version: 1.14.0) was conducted. False Discovery Rate (FDR) was calculated by Benjamini-Hochberg procedure (false_discovery_control from SciPy, version: 1.14.0).

### The time-course analysis of ARG dissemination

The ‘CollectionDate_BIOSAMPLE’ column from PLSDB metadata was manually inspected to determine the year each plasmid was reported. When multiple years were listed (e.g., 1950/1955), the older year was used by default. However, when the older year was before 1900, the more recent year was used (e.g., 1800/2014 or 1900/1967). Plasmids lacking values in the “CollectionDate_BIOSAMPLE” column or with unclear date annotations were excluded from time-course analyses. Threshold Year was defined as the year at which the number of detections for each plasmid type reached the first quartile of their respective detection counts as of 2021. It was calculated for each plasmid type, excluding Both, using all ARGs that were encoded by more than 100 plasmids (n = 48). To compare the distribution of Threshold Year values, the Wilcoxon rank-sum test (wilcoxon from SciPy, version: 1.14.0) was conducted. The False Discovery Rate (FDR) was calculated using the Benjamini-Hochberg procedure (false_discovery_control from SciPy, version: 1.14.0).

## RESULTS

### *h-ns* and *psiB* characterize plasmids with broad and narrow host ranges, respectively

To elucidate the mechanisms by which plasmids acquire either broad-host-range (BHR) or narrow-host-range (NHR) characteristics, we conducted a gene enrichment analysis between BHR and NHR conjugative and mobilizable plasmids (Fig. 1a, b). BHR plasmids are defined as those capable of transferring across multiple bacterial taxa (Orders), whereas NHR plasmids are limited to transfer within a single order or a more restricted taxonomic group. We focused our analysis on 943 conjugative and mobilizable plasmids derived from Enterobacterales (701 NHR and 242 BHR plasmids), which account for a major fraction of plasmids with annotated host ranges and include clinically important lineages such as Klebsiella pneumoniae. Given the high conjugation frequency in the enteric environment where Enterobacterales predominantly reside (40), this group provides an ideal system for investigating the evolution of conjugation-based survival strategies.

**Fig. 1:**
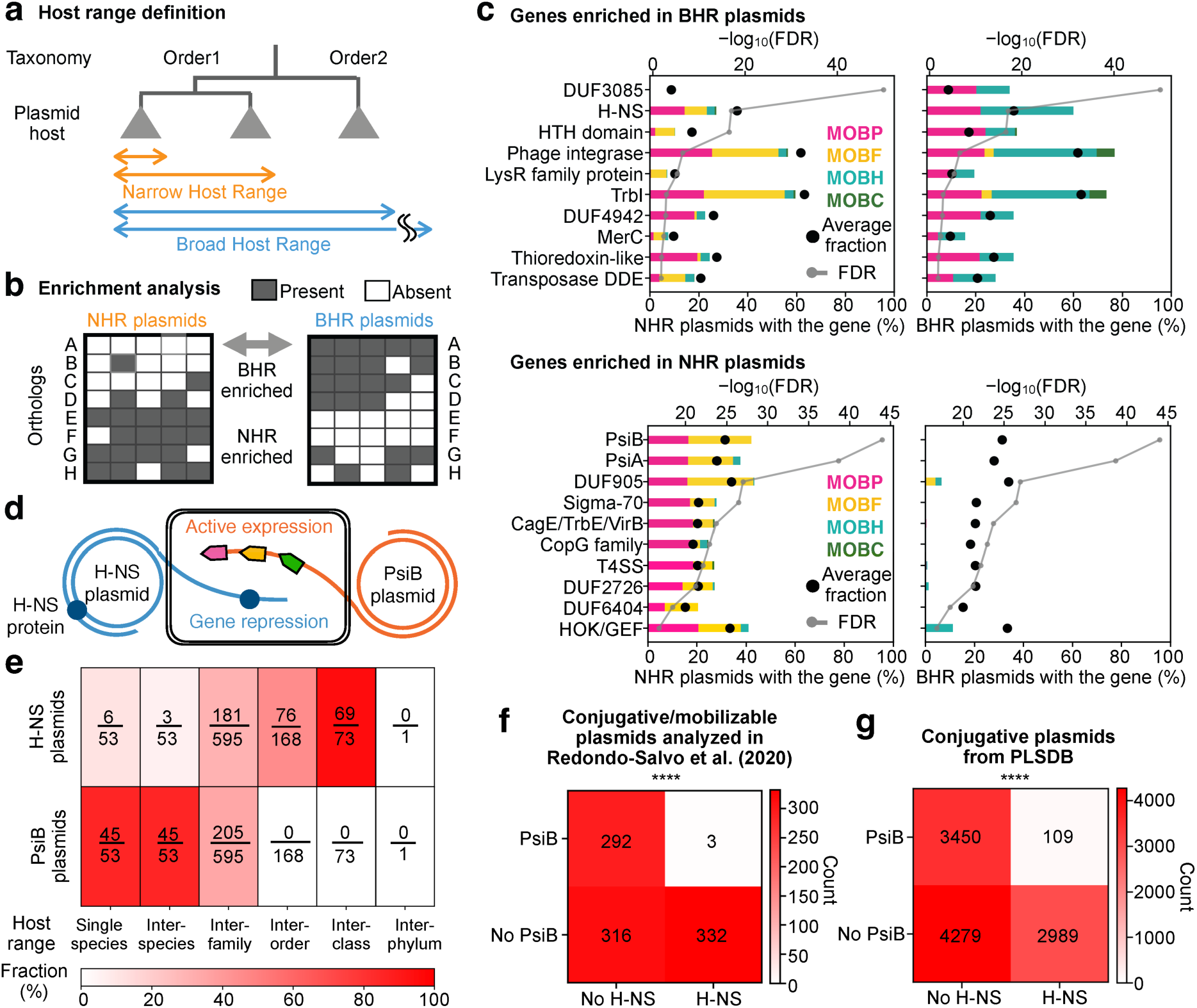
*h-ns* and *psiB* exhibit mutually exclusive distributions and are characteristic of BHR and NHR plasmids, respectively. **a,** Schematic diagram of the host range definition used in this study. **b,** Schematic diagram of the enrichment analysis. Orthologs A–D are enriched in BHR plasmids, whereas E–H are enriched in NHR plasmids. Grey squares indicate presence of orthologs in each plasmid; white squares indicate absence. **c,** Results of the enrichment analysis. Genes were ranked in ascending order of *p*-values from the enrichment analysis among those present in at least 10% of plasmids within each host range and distributed across multiple MOB types (see MATERIAL AND METHODS). Bar colors correspond to the MOB types of the plasmids encoding each gene, while bar heights represent the proportion of plasmids within each host range that carry the gene. Black circles indicate the proportion of plasmids that contain the gene among all analyzed plasmids. For each hit, False Discovery Rate (FDR) was calculated and -log_10_(FDR) of was shown in Gray line. **d,** Schematic representation of the survival strategies employed by H-NS plasmids and PsiB plasmids, respectively. **e,** Proportion of plasmids encoding *h-ns* or *psiB* across different host range categories. “Single species” plasmids have the narrowest host range, being restricted to a single bacterial species, whereas “Inter-phylum” plasmids have the broadest host range, spanning multiple bacterial phyla. **f, g,** Contingency tables showing the distribution of *h-ns* and *psiB* across plasmids. **f,** 943 conjugative/mobilizable plasmids from Enterobacterales annotated by Redondo-Salvo et al. (2020). **g,** 10, 827 conjugative plasmids from PLSDB. Rows indicate the presence or absence of *psiB*, and columns indicate the presence or absence of *h-ns*. The numbers within the boxes indicate the count of plasmids in each category, with color intensity corresponding to plasmid counts. Significance is shown with asterisks (\*\*\*\**p* < 0.0001, Chi-squared test)

Distinct patterns of gene enrichment were observed between BHR and NHR plasmids (Fig. 1c). Expectedly, BHR plasmids were enriched with genes associated with HGT and conjugations. For example, they were enriched with genes central to the integron system, such as integrase (PF00589.25), which facilitates the capture and expression of foreign genes (41, 42). Other examples include *trbI* (PF03743.17), a protein that increases mating efficiency by up to 20-fold during conjugative transfer (43, 44). We further observed that *h-ns* (PF00816.24), a global transcriptional regulator that modulates gene expression and chromosomal structure (45), had the second-lowest p-value in our enrichment analysis, following only the uncharacterized gene DUF3085 (PF11284.11). *h-ns* was both enriched and highly prevalent in BHR plasmids, being encoded in approximately 60% of them, whereas it is found in less than 30% of NHR plasmids. plasmid-encoded *h-ns* not only actively represses the expression of plasmid-borne genes to mitigate the metabolic burden imposed on the host (46, 47) but also to regulate both integrase activity and conjugation frequency (48, 49). Considering the profound impact of *h-ns* on both plasmid gene expression and other machineries including integron and conjugation, the high prevalence of *h-ns* in BHR plasmids suggests that they rely on a lifecycle supported by *h-ns*-mediated transcriptional regulation.

On the other hand, NHR plasmids were associated with genes such as *psiB* (PF06290.14) and *psiA* (PF06952.14). Both *psiB* and *psiA* are known to inhibit the SOS response, particularly during plasmid conjugative transfer. Especially, *psiB* is a well-studied gene, which is known to function independently (50). Additionally, DUF905 (PF06006.15) is often encoded near these genes, suggesting potential functional relationships with them (17, 18, 51–54). Since the SOS response itself does not necessarily affect conjugation, the inhibition of the SOS response by plasmids is thought to serve as a mechanism to avoid unnecessary metabolic costs for the host (52, 55). The inhibition of the SOS response by *psiB* requires a specific protein-protein interaction with the host factor *recA*. Consequently, *psiB* is known to have a restricted range of susceptible bacterial hosts. (17, 52). Therefore, it is reasonable to assume that plasmids utilizing the SOS inhibition mechanism tend to be NHR.

Taken together, these findings suggest that BHR plasmids employ a “stealth strategy” by encoding *h-ns*, which reduces the fitness cost imposed on their hosts while also regulating machineries important for their lifecycle (Fig. 1d). In contrast, while NHR plasmids may also mitigate additional metabolic costs by suppressing the SOS response, this mechanism is entirely dependent on the transcription of their own genes. Moreover, among the genes enriched in NHR plasmids, no global transcriptional repressors or genes typically associated with “stealth” strategies—such as H-NS—were identified. Based on these findings, we hypothesized that NHR plasmids adopt an “aggressive strategy, “ prioritizing their own maintenance and propagation through active transcription, probably by inhibiting machinery in its host (Fig. 1d). In the following analyses, *h-ns* and *psiB* were selected as marker genes of these distinct strategies, and plasmids encoding either gene were referred to as H-NS plasmids or PsiB plasmids, respectively.

We found that the proportion of plasmids encoding *h-ns* increases as the host range broadens, while *psiB* exhibits the opposite trend (Fig. 1e), which is consistent with previous findings that associate *h-ns* and *psiB* with some BHR and NHR plasmids, respectively (17, 56). Moreover, these genes were distributed almost mutually exclusively (Fig. 1f). This result supports the hypothesis that NHR and BHR plasmids adopt to their respective host ranges based on mutually exclusive survival strategies, with *psiB* and *h-ns* serving as (at least, ones of the) critical genes for these strategies. Jaccard index between the distribution of *h-ns* and *psiB* was 3.69 × 10⁻^3^, which was the 560th smallest out of 12, 882 randomly sampled gene pairs analyzed (Supplementary Fig. 1). Such an exclusive distribution was robustly observed in a comprehensive analysis of all conjugative plasmids (10, 827 sequences) available in the PLSDB database (28, 29) (Fig. 1g).

### H-NS and PsiB plasmids distributed across diverse plasmid taxa and have undergone distinct adaptations

The mutually exclusive distribution of these genes could be an artefact of phylogenetically (taxonomically) biased distribution across diverse plasmid taxonomy. To rule out this possibility, we next examined the taxonomic group of plasmids carrying these genes. Inferring accurate phylogenetics of plasmid lineages is challenging due to frequent recombination events with host genomes and other MGEs, as well as the absence of essential genes. However, MOB typing, which classifies plasmids based on the sequence of relaxase—a protein essential for conjugative transfer and present in all mobilizable and conjugative plasmids—provides a method for phylogenetic classification to a certain extent (57–59). The sequences of relaxase with different MOB types differ substantially and rarely coexist on a single plasmid. Moreover, relaxase evolves very slowly compared to other features such as plasmid size or other encoded genes(58). These facts make it possible to assume that plasmids belonging to different MOB types are phylogenetically more distant from each other than those within the same MOB type.

Using 10, 827 conjugative plasmids from PLSDB, we analyzed the MOB-type compositions of plasmids in our dataset (Fig. 2a). H-NS and PsiB plasmids were distributed across diverse MOB types, which indicates that *h-ns* and *psiB* have been independently acquired by various clades. Notably, certain MOB types, such as MOBP and MOBF, contained both H-NS plasmids and PsiB plasmids, and their mutually exclusive distribution was statistically significant (Fig. 2a). The fact that a single taxonomic group of plasmids contains both types of plasmids, yet the number of plasmids with both genes remains very small, implies that this exclusivity cannot be solely explained by phylogenetic bias. Instead, it suggests a nontrivial evolutionary pattern driven by distinct selective pressures.

**Fig. 2:**
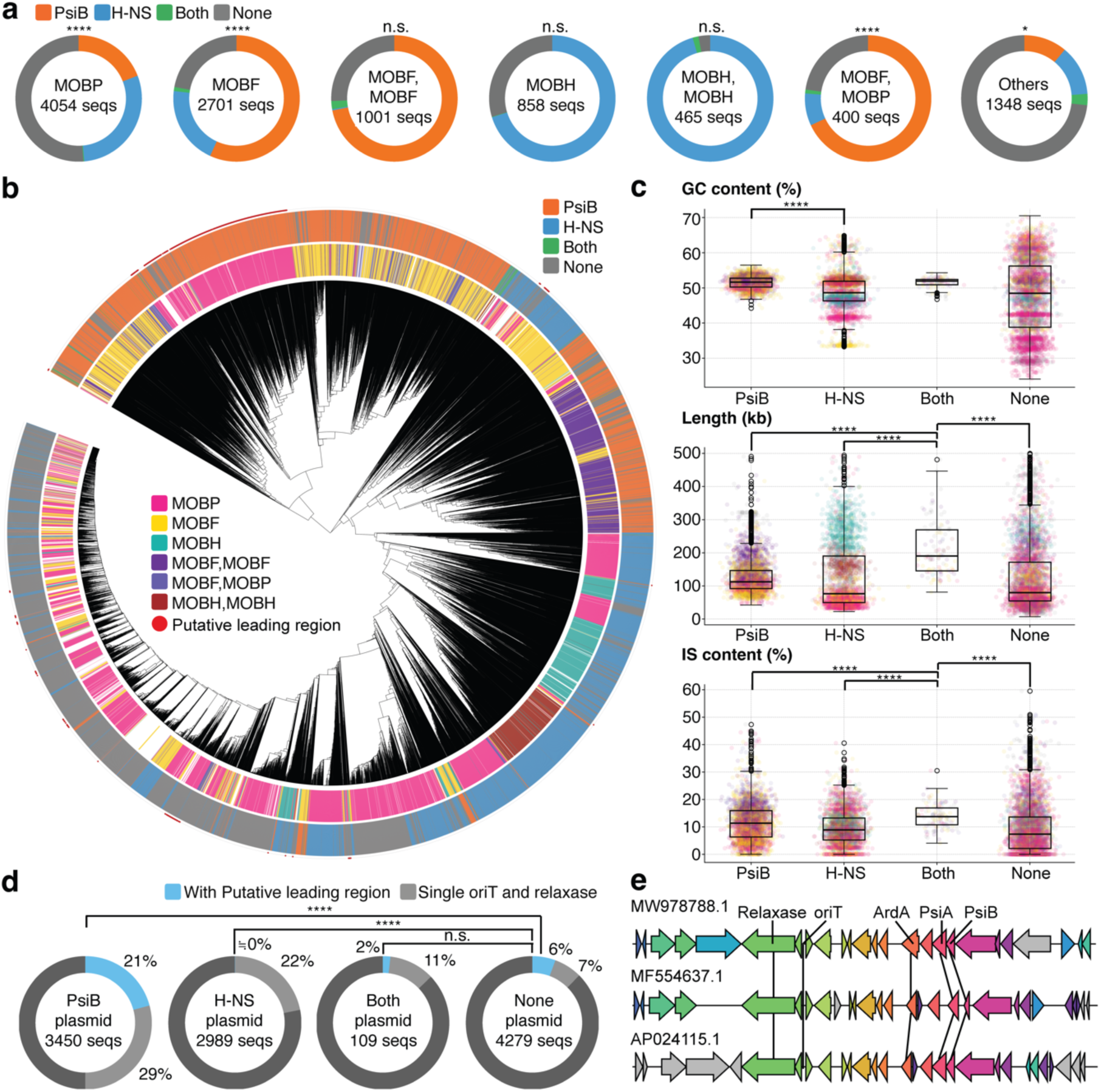
H-NS and PsiB plasmids originate from diverse plasmid clades and exhibit distinct adaptations. **a,** Proportion of H-NS and PsiB plasmids for each MOB type among all analyzed conjugative plasmids (10, 827 sequences). The six most abundant MOB types, as well as the sum of all remaining MOB types, are shown. Significance is shown with asterisks (\**p* < 0.05; \*\*\*\**p* < 0.0001, Chi-squared test). **b,** Dendrogram based on the Average Nucleotide Identity (ANI) of all analyzed conjugative plasmids. The outer ring indicates plasmid types. The color of the inner ring represents the MOB type of each plasmid. Asterisks mark plasmids annotated with putative leading regions. **c,** Characteristics of each plasmid type. Dot colors correspond to plasmid MOB types, with color assignments matching those in Fig. 2b. Significance is shown with asterisks (****FDR < 0.0001, Mann-Whitney U test). **d,** Proportion of H-NS and PsiB plasmids carrying putative leading regions. Putative leading regions were defined as plasmids encoding one relaxase and one oriT within 1, 000 bp of each other. Significance is shown with asterisks (****FDR < 0.0001, Chi-squared test). **e,** Example of a putative leading region. Relaxase and oriT(origin of Transfer) were annotated using MOB-suite, while other genes were annotated using HMMER (see MATERIAL AND METHODS).

Furthermore, on a dendrogram constructed based on the average nucleotide identity (ANI) of all plasmids, both plasmid types showed a patchy distribution (Fig. 2b). This observation further indicates that the acquisition of PsiB and H-NS, along with the evolution of survival strategies based on these genes, has occurred repeatedly across different clades.

Next, to investigate the adaptive strategies of H-NS and PsiB plasmids, we examined the characteristics of plasmids carrying H-NS or PsiB, as well as those carrying neither or both genes (“None” and “Both”). H-NS plasmids exhibited notably lower GC contents compared to PsiB plasmids (Mann-Whitney U test, FDR = 4.44 × 10^−196^) (Fig. 2c). Given that *h-ns* is a transcriptional repressor with a preference for AT-rich DNA (60, 61), this suggests sequence-level adaptation for transcriptional regulation centered around *h-ns*. Furthermore, each plasmid type exhibited distinct size distributions. “Both” plasmids (median = 190 kbp) had a significantly larger size distribution, with a median 1.69 times greater than that of the next largest group, PsiB plasmids (median = 112 kbp) (Fig. 2c). The scarcity of ‘Both’ plasmids, along with the fact that they are larger and contain more genes per plasmid compared to other groups, suggests that these plasmids may have acquired sequences from multiple plasmids through recombination or adopted a unique survival strategy that promotes the accumulation of diverse genes. GC content and plasmid length both showed relatively distinct distributions across different MOB families, indicating that mobility type contributes to the overall architecture of plasmids. However, variation in plasmid size could not be fully explained by MOB composition alone. These size differences also mirrored the proportion of insertion sequences within their nucleotide sequences, suggesting that IS elements act as additional drivers of plasmid size variation beyond what is determined by MOB family constraints (Fig. 2c).

As another sequence feature that reflects the distinct survival strategies of conjugative plasmids, we further analyzed the presence of leading regions. A leading region is a genetic structure located near the origin of transfer (oriT) that enhances plasmid survival by enabling the early expression of genes immediately upon entry into the recipient cell. These regions often encode proteins that inhibit host responses such as various defense systems and SOS response, thereby increasing the likelihood of successful plasmid establishment during conjugation (51, 54, 62). We therefore hypothesized that plasmids employing an aggressive strategy—characterized by active transcription—would be more likely to possess a leading region than those relying on a stealth strategy. Indeed, our analysis confirmed this hypothesis: approximately 21% of PsiB plasmids contained putative leading regions, whereas a putative leading region was observed in only one H-NS plasmid. Compared to None plasmids, PsiB plasmids showed a significant enrichment of leading regions (Chi-squared test, FDR = 2.26 × 10^−89^). Conversely, H-NS plasmids exhibited a significant depletion of leading regions (Chi-squared test, FDR = 2.59 × 10^−38^) (Fig. 2d). Among the detected leading regions, several characteristic genes were frequently observed in addition to *psiB*, such as *ardA* (an anti-restriction gene) (Fig. 2e). This observation suggests that *psiB* in aggressive plasmids is often utilized as part of the leading region system. Collectively, these data reinforce the idea that PsiB and H-NS plasmids employ two opposing survival strategies.

### Both H-NS plasmids and PsiB plasmids are significant contributors to the dissemination of ARGs

The aforementioned analyses suggested that H-NS and PsiB plasmids have undergone distinct evolutionary adaptations and have employed mutually exclusive survival strategies. To explore how these strategies influence the spread of ARGs, we analyzed ARG accumulation across these plasmid types (Fig. 3a). Interestingly, PsiB plasmids, H-NS plasmids, and Both plasmids accumulated more ARGs than None plasmids (Mann-Whitney U test, FDR = 4.13 × 10⁻⁷⁴, 2.88 × 10^−288^, and 7.22 × 10⁻²⁶, respectively). H-NS plasmids accumulated significantly more ARGs than PsiB plasmids (Mann-Whitney U test, FDR = 2.67 × 10⁻^109^), the median number of ARGs normalized by plasmid size being 2.87 times higher in H-NS plasmids that in PsiB plasmids. These results suggest that, in the context of ARG dissemination by conjugative plasmids, plasmids carrying *h-ns* and/or *psiB* play a significantly more important role compared to those lacking these genes. Since integrons are known to play a critical role in the acquisition of ARGs in MGEs (63), we examined the proportion of integron regions in the sequences of each plasmid belonging to each plasmid type (Fig. 3b). As expected, PsiB plasmids, H-NS plasmids, and Both plasmids contained a higher proportion of plasmids with integron regions compared to None plasmids (Mann-Whitney U test, FDR = 3.14 × 10^−61^, 1.09 × 10^−144^, and 8.47 × 10^−43^, respectively). Additionally, H-NS plasmids contained more plasmids with integron regions than PsiB plasmids (Mann-Whitney U test, FDR = 6.94 × 10⁻²⁴). This pattern aligns with the trends observed for the number of ARGs per 100 kbp.

**Fig. 3:**
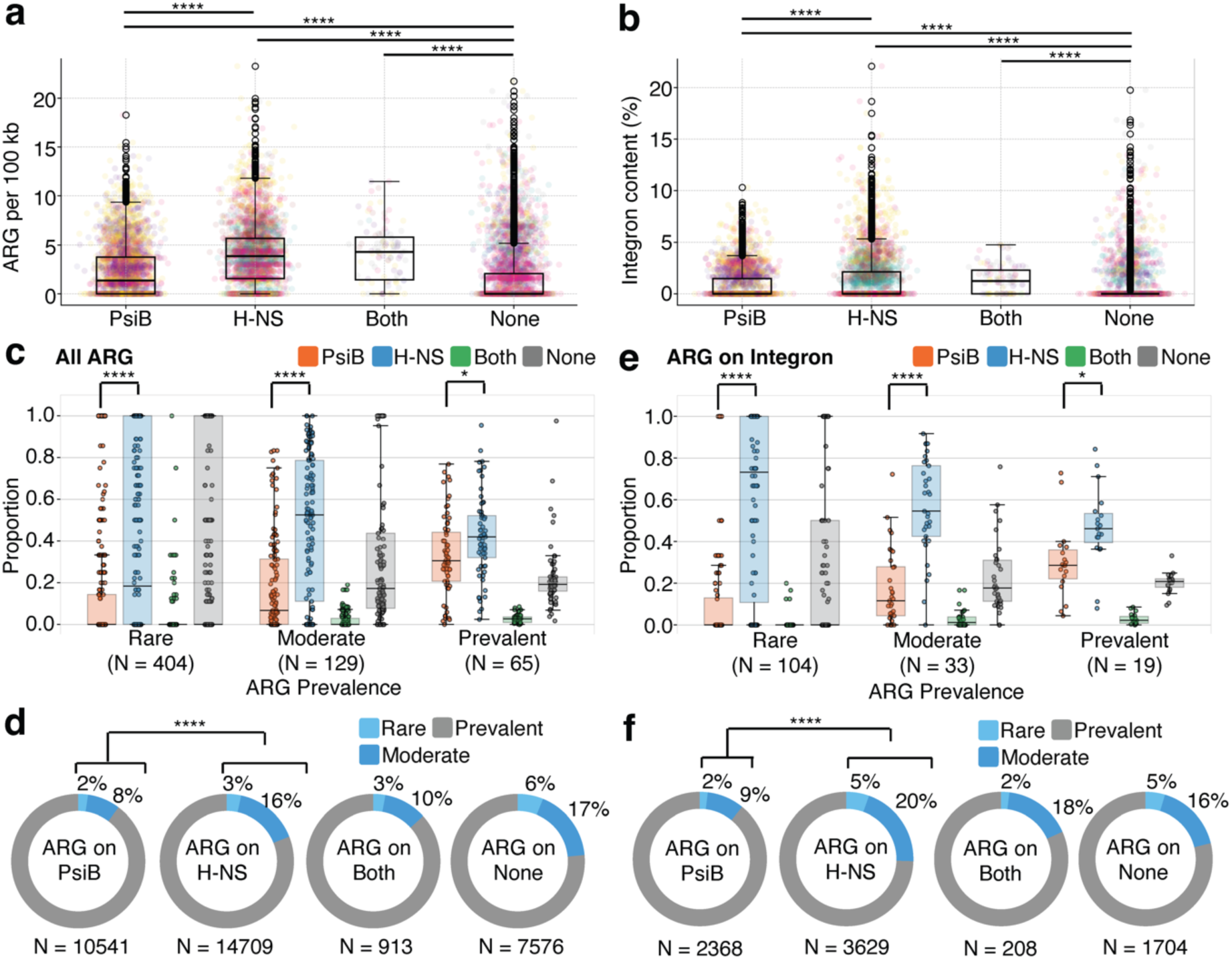
Conjugative plasmids carrying *h-ns* or *psiB* are major contributors to ARG dissemination. **a,** Number of ARGs per 100 kb in each plasmid type. Dot colors correspond to plasmid MOB types, with color assignments matching those in Fig. 2b. Significance is shown with asterisks (****FDR < 0.0001, Mann-Whitney U test). **c, e,** Proportion of plasmids carrying ARGs by plasmid type and ARG prevalence. ARGs were categorized based on the number of plasmids encoding them: ARGs present in fewer than 10 plasmids were classified as ‘Rare’, those found in 10 to 99 plasmids as ‘Moderate’, and those present in 100 or more plasmids as ‘Prevalent’. The numbers of unique ARGs within each prevalence category are shown in the plot. Significance is shown with asterisks (*FDR < 0.05; ****FDR < 0.0001, Wilcoxon rank-sum test). **d, f,** Proportions of ARGs in each prevalence category detected in each plasmid type. The total numbers of ARGs detected in each plasmid type are shown in the plot. Significance is shown with asterisks (****FDR < 0.0001, Chi-squared test). **c, d,** Analysis on all detected ARGs. **e, f,** Analysis on ARGs located within integron regions.

We then examined the characteristics of ARGs found on each plasmid type. The detected ARGs in the dataset were classified into three categories based on the number of plasmids encoding each ARG. ARGs encoded by fewer than 10 plasmids were classified as ‘Rare’, those found in 10 to 99 plasmids as ‘Moderate’, and those present in 100 or more plasmids as ‘Prevalent’. For each prevalence category, we plotted the proportion of plasmid types encoding each ARG. The results indicate that although H-NS plasmids generally carry ARGs more frequently than PsiB plasmids, the difference in ARG carriage frequency between the two plasmid types becomes smaller for prevalent ARGs (Fig. 3c). Furthermore, among the ARGs detected in H-NS plasmids, the proportion classified as ‘Rare’ or ‘Moderate’ was higher than in PsiB plasmids (Chi-squared test, FDR = 8.31 × 10⁻⁸¹) (Fig. 3d). These data suggest that rare ARGs, which have not yet widely spread within a population, are preferentially acquired by H-NS plasmids. The trend became even more pronounced when only the ARGs located within integron regions (Fig. 3e, f) were analyzed. These results suggest that H-NS plasmids tend to capture ARGs that have not yet been widely spread in the population, possibly facilitated by integrons.

### H-NS plasmids acquire ARGs earlier than other conjugative plasmids

Our findings so far suggest that H-NS plasmids contribute to the acquisition of rare or emerging ARGs. If this hypothesis is correct, the acquisition of ARGs by H-NS plasmids is expected to precede their acquisition by PsiB plasmids. To test this, we analyzed all ARGs that were encoded by more than 100 plasmids (n = 48), focusing in particular on the temporal distribution of plasmids carrying the same ARG. For illustrative purposes, we first highlight the top 10 most frequently detected ARGs in the dataset. For all of these top 10 ARGs except *mph(A)* and *aadA1*, a sharp increase in the detection of ARGs on H-NS plasmids preceded the corresponding increase on PsiB plasmids (Fig. 4a, Supplementary Fig. 3). These findings support a model in which H-NS plasmids, which tend to exhibit a broad host range, act as hubs for the early acquisition and dissemination of ARGs by conjugative plasmids. They likely achieve this by capturing ARGs from diverse environments and subsequently distributing them within the plasmid community (Fig. 4b). To assess whether this trend holds across a broader range of ARGs, we compared the distribution of “threshold years“—the year at which each plasmid type reached the first quartile of its cumulative detection count as of 2021—for all 48 ARGs (Fig. 4c). The resulting distribution showed that the acquisition of ARGs by H-NS and PsiB plasmids significantly preceded that by None plasmids (Wilcoxon rank-sum test, FDR = 1.42 × 10⁻⁶ and 1.89 × 10⁻³, respectively), and that acquisition by H-NS plasmids preceded that by PsiB plasmids (FDR = 1.70 × 10⁻⁴) (Fig. 4d). This trend—namely, the precedence of H-NS and PsiB plasmids over None plasmids, as well as the precedence of H-NS plasmids over PsiB plasmids—was consistently observed when calculating the differences in Threshold Years across plasmid types for each ARG (Fig. 4e). It indicates that, both for individual ARGs and as an overall trend, the timing of ARG acquisition on H-NS plasmids precedes that on PsiB plasmids. Moreover, throughout the entire period, the number of PsiB plasmids reported in PLSDB was consistently similar to that of H-NS plasmids, with PsiB plasmids being slightly more prevalent, confirming that the observed tendency for ARG acquisition by H-NS plasmids to precede that of PsiB plasmids is not attributable to sampling bias (Fig. 4f).

**Fig. 4:**
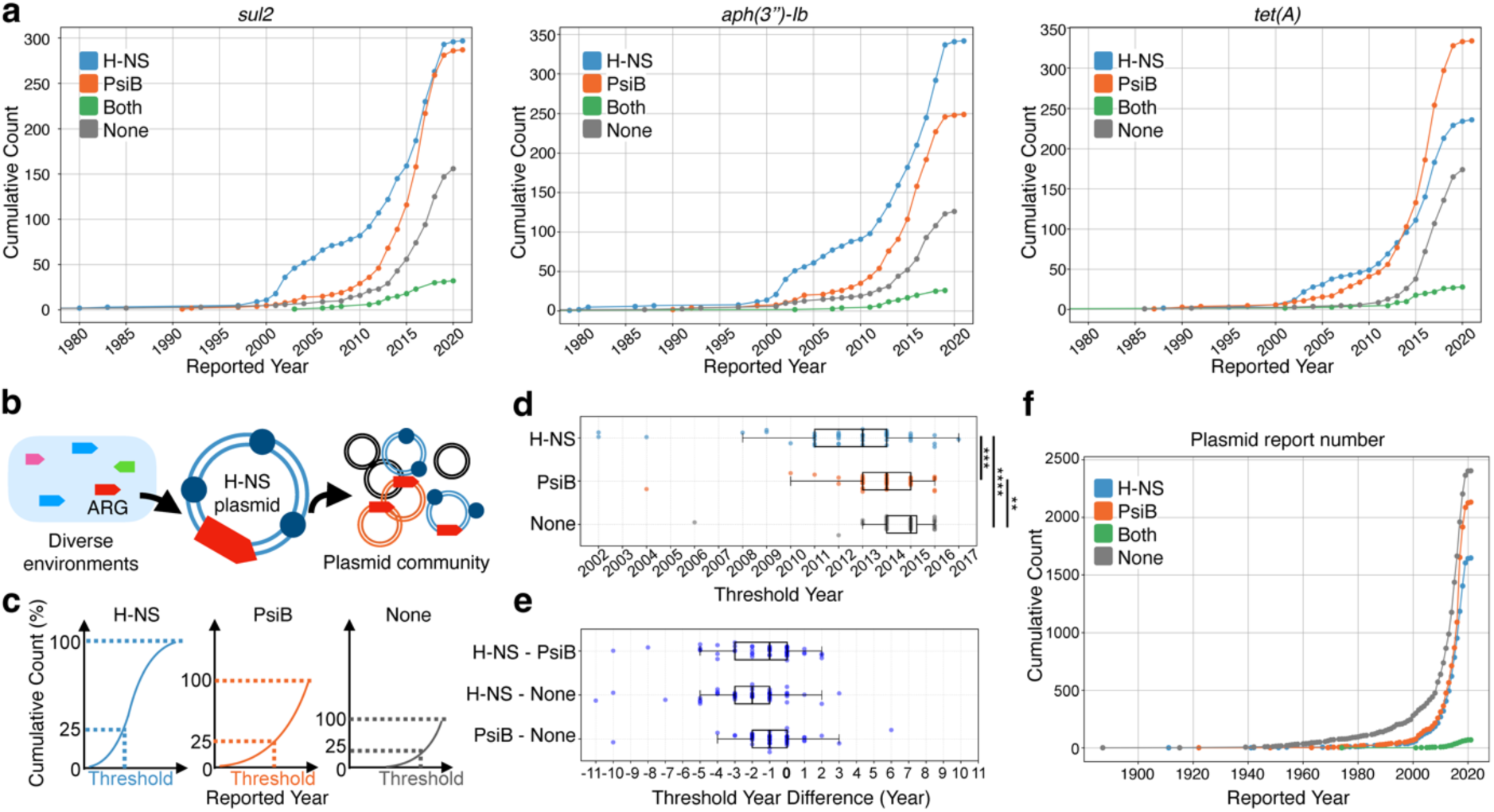
ARG acquisition by H-NS plasmids precedes that by PsiB plasmids. **a,** Cumulative frequency curves illustrating the temporal increase in the number of plasmid records encoding *sul2*, *aph(3’’)-Ib*, and *tet(A)* based on the collection dates registered in the PLSDB database (see MATERIAL AND METHODS). **b,** Schematic diagram of the model of ARG spread mediated by H-NS plasmids. **c,** Schematic diagram illustrating the calculation method of the Threshold Year. **d,** Distribution of Threshold Year values for all ARGs that were encoded by more than 100 plasmids (n = 48). Significance is shown with asterisks (**FDR < 0.01; ***FDR <0.001; ****FDR < 0.0001, Wilcoxon rank-sum test). **e,** Distribution of the differences in Threshold Year values between plasmid types for the same ARG, calculated for all ARGs encoded by more than 100 plasmids (n = 48). **f,** Cumulative frequency curves showing the numbers of reported plasmid records by collection year for each plasmid type.

## DISCUSSION

This study reveals that among conjugative plasmids, distinct survival strategies are closely linked to host range and contribute to the dissemination of antimicrobial resistance genes (ARGs). *H-NS* and *PsiB*—two key regulatory proteins independently acquired across diverse plasmid lineages—characterize plasmids with broad and narrow host ranges, respectively (Fig. 1e, Fig. 2b). These plasmids adopt contrasting survival strategies: a “stealth” strategy in H-NS plasmids, which minimizes host fitness cost, and an “aggressive” strategy in PsiB plasmids, which suppresses host stress responses. These results extend previous studies that have focused on broad genomic features affecting plasmid host range—such as restriction site avoidance, methyltransferase copy number, and replication origin structure (25, 26). By integrating recent advances in plasmid host range prediction (20) with our own insights into specific regulatory genes and the survival strategies they encode, our study identifies functional determinants of host range at the gene and strategy level—linking molecular function with ecological adaptation. These results align with recent work showing that distinct survival strategies adopted by different MGEs shape their accessory gene repertoires (64), and extend this principle to host range–dependent strategies in conjugative plasmids. We also found that leading regions—a genetic mechanism that enhances plasmid survival by promoting the early expression of genes that inhibit host response (51, 54, 62)— were markedly more prevalent in PsiB plasmids (Fig. 2d). This supports the interpretation that leading regions are an integral component of the aggressive strategy employed by these plasmids.

Importantly, both H-NS and PsiB plasmids harbored significantly more ARGs than plasmids lacking either gene, suggesting that both the stealth and aggressive strategies promote ARG dissemination (Fig. 3a). However, they do so in distinct ways: H-NS plasmids, with their broad host range, appear to act as early hubs that acquire novel ARGs from diverse environments, whereas PsiB plasmids contribute to the amplification of already prevalent ARGs within narrower ecological niches. This “stealth-first” model recapitulates historical ARG dissemination patterns (Fig. 4a). We further observed that integrons may contribute to the acquisition of ARGs by H-NS plasmids. Although *h-ns* is known to enhance integrase expression while repressing the transcription of cassette genes (49), it has remained unclear whether its overall effect on integron function is beneficial or detrimental. Our results suggest that, at least in the context of ARG acquisition, the regulatory characteristics of *h-ns*—or other features associated with H-NS plasmids—are compatible with, and may even promote integron-mediated gene capture.

Evolutionary predictability has been demonstrated through the recurrent emergence of beneficial mutations across lineages (65), and forecasting the evolution of antibiotic resistance remains an urgent challenge (66). Mobile genetic elements (MGEs), which mediate horizontal gene transfer, are central to the spread of resistance genes (67). While ARG acquisition is beneficial under antibiotic pressure, it imposes a cost in its absence— making its dynamics context-dependent and difficult to model. Recent approaches incorporate gene transfer networks and genome-wide acquisition/loss patterns to better understand HGT-prone ARGs (67, 68). Complementing this line of research, our study provides insights from the MGE perspective by identifying characteristic features of plasmids that are prone to acquiring and disseminating ARGs. Here, we identify a generalizable signature of ARG dissemination: ARGs consistently emerge first on “stealth” MGEs before appearing on others. This temporal pattern offers a generalizable framework for inferring the stages of ARG dissemination, with stealth-exclusive ARGs representing early-stage variants and those found across multiple MGE types reflecting broader epidemiological spread. Together, these findings lay a foundation for predictive frameworks that integrate MGE biology into the surveillance and control of antimicrobial resistance.

## DATA AVAILABILITY

The data underlying this article are available within the article and its online supplementary material. Additional supporting data are available in the Figshare repository at: https://figshare.com/s/49fad8b5e286b216cf16. (Private link)

## Supplementary Data statement

Supplementary Data are available at NAR online.

## AUTHOR CONTRIBUTIONS

Conceptualization: R.O. and N.K., Methodology: R.O., N.K., and C.F., Investigation: R.O., Visualization: R.O., Supervision: N.K., Writing-original draft: R.O. and N.K., Writing-review and editing: R.O., N.K., Y.N., and C.F.

## ACKNOWLEDGEMENTS

We thank all members of the Furusawa and Iwasaki labs for their valuable discussions on the content of this paper, especially W. Iwasaki for his feedback on our work. We also thank B. Cress for his insight and support. This work was supported by the Japan Society for the Promotion of Science (KAKENHI Grant number 22H04925) and the Japan Science and Technology Agency (GteX Program JPMJGX23B2).

## REFERENCES

1. Treangen, T.J. and Rocha, E.P.C. (2011) Horizontal transfer, not duplication, drives the expansion of protein families in prokaryotes. PLoS Genet., 7, e1001284.

2. Puigbò, P., Lobkovsky, A.E., Kristensen, D.M., Wolf, Y.I. and Koonin, E.V. (2014) Genomes in turmoil: quantification of genome dynamics in prokaryote supergenomes. BMC Biol., 12, 66.

3. Smillie, C.S., Smith, M.B., Friedman, J., Cordero, O.X., David, L.A. and Alm, E.J. (2011) Ecology drives a global network of gene exchange connecting the human microbiome. Nature, 480, 241–244.

4. Andam, C.P. and Gogarten, J.P. (2011) Biased gene transfer in microbial evolution. Nat. Rev. Microbiol., 9, 543–555.

5. Soucy, S.M., Huang, J. and Gogarten, J.P. (2015) Horizontal gene transfer: building the web of life. Nat. Rev. Genet., 16, 472–482.

6. Jain, R., Rivera, M.C. and Lake, J.A. (1999) Horizontal gene transfer among genomes: The complexity hypothesis. Proc. Natl. Acad. Sci. U. S. A., 96, 3801–3806.

7. Burch, C.L., Romanchuk, A., Kelly, M., Wu, Y. and Jones, C.D. (2023) Empirical evidence that complexity limits horizontal gene transfer. Genome Biol. Evol., 15.

8. Kogay, R., Wolf, Y.I. and Koonin, E.V. (2024) Defense systems and horizontal gene transfer in bacteria. bioRxiv, 10.1101/2024.02.09.579689.

9. Neil, K., Allard, N., Grenier, F., Burrus, V. and Rodrigue, S. (2020) Highly efficient gene transfer in the mouse gut microbiota is enabled by the Incl2 conjugative plasmid TP114. Commun. Biol., 3, 523.

10. Rubin, B.E., Diamond, S., Cress, B.F., Crits-Christoph, A., Lou, Y.C., Borges, A.L., Shivram, H., He, C., Xu, M., Zhou, Z., et al. (2022) Species- and site-specific genome editing in complex bacterial communities. Nat Microbiol, 7, 34–47.

11. Murray, C.J.L., Ikuta, K.S., Sharara, F., Swetschinski, L., Robles Aguilar, G., Gray, A., Han, C., Bisignano, C., Rao, P., Wool, E., et al. (2022) Global burden of bacterial antimicrobial resistance in 2019: a systematic analysis. Lancet, 399, 629–655.

12. O’Neill, J. (2016) Tackling drug-resistant infections globally: final report and recommendations.

13. World Bank (2017) Drug-Resistant Infections: A threat to our economic future World Bank, Washington, DC.

14. Castañeda-Barba, S., Top, E.M. and Stalder, T. (2024) Plasmids, a molecular cornerstone of antimicrobial resistance in the One Health era. Nat. Rev. Microbiol., 22, 18–32.

15. Che, Y., Yang, Y., Xu, X., Břinda, K., Polz, M.F., Hanage, W.P. and Zhang, T. (2021) Conjugative plasmids interact with insertion sequences to shape the horizontal transfer of antimicrobial resistance genes. Proc. Natl. Acad. Sci. U. S. A., 118.

16. Wang, X., Zhang, H., Long, X., Xu, X., Ren, H., Mao, D., Alvarez, P.J.J. and Luo, Y. (2023) Global increase of antibiotic resistance genes in conjugative plasmids. Microbiol. Spectr.

17. Baharoglu, Z., Bikard, D. and Mazel, D. (2010) Conjugative DNA transfer induces the bacterial SOS response and promotes antibiotic resistance development through integron activation. PLoS Genet., 6, e1001165.

18. Pons, M.C., Praud, K., Da Re, S., Cloeckaert, A. and Doublet, B. (2023) Conjugative IncC Plasmid entry triggers the SOS response and promotes effective transfer of the integrative antibiotic resistance element SGI1. Microbiol. Spectr., 11, e0220122.

19. Klümper, U., Riber, L., Dechesne, A., Sannazzarro, A., Hansen, L.H., Sørensen, S.J. and Smets, B.F. (2015) Broad host range plasmids can invade an unexpectedly diverse fraction of a soil bacterial community. ISME J., 9, 934–945.

20. Redondo-Salvo, S., Fernández-López, R., Ruiz, R., Vielva, L., de Toro, M., Rocha, E.P.C., Garcillán-Barcia, M.P. and de la Cruz, F. (2020) Pathways for horizontal gene transfer in bacteria revealed by a global map of their plasmids. Nat. Commun., 11, 3602.

21. Heinemann, J.A. and Sprague, G.F., Jr (1989) Bacterial conjugative plasmids mobilize DNA transfer between bacteria and yeast. Nature, 340, 205–209.

22. Waters, V.L. (2001) Conjugation between bacterial and mammalian cells. Nat. Genet., 29, 375–376.

23. Bertozzi Silva, J., Storms, Z. and Sauvageau, D. (2016) Host receptors for bacteriophage adsorption. FEMS Microbiol. Lett., 363, fnw002.

24. Forster, S.C., Liu, J., Kumar, N., Gulliver, E.L., Gould, J.A., Escobar-Zepeda, A., Mkandawire, T., Pike, L.J., Shao, Y., Stares, M.D., et al. (2022) Strain-level characterization of broad host range mobile genetic elements transferring antibiotic resistance from the human microbiome. Nat. Commun., 13, 1445.

25. Shaw, L.P., Rocha, E.P.C. and MacLean, R.C. (2023) Restriction-modification systems have shaped the evolution and distribution of plasmids across bacteria. Nucleic Acids Res., 51, 6806–6818.

26. Jain, A. and Srivastava, P. (2013) Broad host range plasmids. FEMS Microbiol. Lett., 348, 87–96.

27. Pruitt, K.D., Tatusova, T. and Maglott, D.R. (2007) NCBI reference sequences (RefSeq): a curated non-redundant sequence database of genomes, transcripts and proteins. Nucleic Acids Res., 35, D61–5.

28. Galata, V., Fehlmann, T., Backes, C. and Keller, A. (2019) PLSDB: a resource of complete bacterial plasmids. Nucleic Acids Res., 47, D195–D202.

29. Schmartz, G.P., Hartung, A., Hirsch, P., Kern, F., Fehlmann, T., Müller, R. and Keller, A. (2022) PLSDB: advancing a comprehensive database of bacterial plasmids. Nucleic Acids Res., 50, D273–D278.

30. Eddy, S.R. (2011) Accelerated profile HMM searches. PLoS Comput. Biol., 7, e1002195.

31. Robertson, J. and Nash, J.H.E. (2018) MOB-suite: software tools for clustering, reconstruction and typing of plasmids from draft assemblies. Microb Genom, 4.

32. Xie, Z. and Tang, H. (2017) ISEScan: automated identification of insertion sequence elements in prokaryotic genomes. Bioinformatics, 33, 3340–3347.

33. Jain, C., Rodriguez-R, L.M., Phillippy, A.M., Konstantinidis, K.T. and Aluru, S. (2018) High throughput ANI analysis of 90K prokaryotic genomes reveals clear species boundaries. Nat. Commun., 9, 5114.

34. moshi ANIclustermap: A tool for drawing ANI clustermap between all-vs-all microbial genomes Github.

35. Letunic, I. and Bork, P. (2024) Interactive Tree of Life (iTOL) v6: recent updates to the phylogenetic tree display and annotation tool. Nucleic Acids Res., 52, W78–W82.

36. Ares-Arroyo, M., Nucci, A. and Rocha, E.P.C. (2024) Identification of novel origins of transfer across bacterial plasmids. bioRxiv, 10.1101/2024.01.30.577996.

37. Gilchrist, C.L.M. and Chooi, Y.-H. (2021) Clinker & clustermap.Js: Automatic generation of gene cluster comparison figures. Bioinformatics, 37, 2473–2475.

38. Feldgarden, M., Brover, V., Gonzalez-Escalona, N., Frye, J.G., Haendiges, J., Haft, D.H., Hoffmann, M., Pettengill, J.B., Prasad, A.B., Tillman, G.E., et al. (2021) AMRFinderPlus and the Reference Gene Catalog facilitate examination of the genomic links among antimicrobial resistance, stress response, and virulence. Sci. Rep., 11, 12728.

39. Néron, B., Littner, E., Haudiquet, M., Perrin, A., Cury, J. and Rocha, E.P.C. (2022) IntegronFinder 2.0: Identification and analysis of integrons across bacteria, with a focus on antibiotic resistance in Klebsiella. Microorganisms, 10, 700.

40. Liu, L., Chen, X., Skogerbø, G., Zhang, P., Chen, R., He, S. and Huang, D.-W. (2012) The human microbiome: a hot spot of microbial horizontal gene transfer. Genomics, 100, 265–270.

41. Mazel, D. (2006) Integrons: agents of bacterial evolution. Nat. Rev. Microbiol., 4, 608–620.

42. Gillings, M.R. (2014) Integrons: past, present, and future. Microbiol. Mol. Biol. Rev.

43. Maneewannakul, S., Maneewannakul, K. and Ippen-Ihler, K. (1992) Characterization, localization, and sequence of F transfer region products: the pilus assembly gene product TraW and a new product, TrbI. J. Bacteriol., 174, 5567–5574.

44. Arutyunov, D., Arenson, B., Manchak, J. and Frost, L.S. (2010) F plasmid TraF and TraH are components of an outer membrane complex involved in conjugation. J. Bacteriol., 192, 1730–1734.

45. Dorman, C.J. (2004) H-NS: a universal regulator for a dynamic genome. Nat. Rev. Microbiol., 2, 391–400.

46. Doyle, M., Fookes, M., Ivens, A., Mangan, M.W., Wain, J. and Dorman, C.J. (2007) An H-NS-like stealth protein aids horizontal DNA transmission in bacteria. Science, 315, 251–252.

47. Shintani, M., Suzuki-Minakuchi, C. and Nojiri, H. (2015) Nucleoid-associated proteins encoded on plasmids: Occurrence and mode of function. Plasmid, 80, 32–44.

48. Forns, N., Baños, R.C., Balsalobre, C., Juárez, A. and Madrid, C. (2005) Temperature-dependent conjugative transfer of R27: role of chromosome- and plasmid-encoded Hha and H-NS proteins. J. Bacteriol., 187, 3950–3959.

49. Cagle, C.A., Shearer, J.E.S. and Summers, A.O. (2011) Regulation of the integrase and cassette promoters of the class 1 integron by nucleoid-associated proteins. Microbiology, 157, 2841–2853.

50. Bailone, A., Bäckman, A., Sommer, S., Célérier, J., Bagdasarian, M.M., Bagdasarian, M. and Devoret, R. (1988) PsiB polypeptide prevents activation of RecA protein in Escherichia coli. Mol. Gen. Genet., 214, 389–395.

51. Althorpe, N.J., Chilley, P.M., Thomas, A.T., Brammar, W.J. and Wilkins, B.M. (1999) Transient transcriptional activation of the Incl1 plasmid anti-restriction gene (ardA) and SOS inhibition gene (psiB) early in conjugating recipient bacteria. Mol. Microbiol., 31, 133–142.

52. Petrova, V., Chitteni-Pattu, S., Drees, J.C., Inman, R.B. and Cox, M.M. (2009) An SOS inhibitor that binds to free RecA protein: the PsiB protein. Mol. Cell, 36, 121–130.

53. Petrova, V., Satyshur, K.A., George, N.P., McCaslin, D., Cox, M.M. and Keck, J.L. (2010) X-ray crystal structure of the bacterial conjugation factor PsiB, a negative regulator of RecA. J. Biol. Chem., 285, 30615–30621.

54. Samuel, B., Mittelman, K., Croitoru, S.Y., Ben Haim, M. and Burstein, D. (2024) Diverse anti-defence systems are encoded in the leading region of plasmids. Nature, 635, 186–192.

55. Alam, M.K., Alhhazmi, A., DeCoteau, J.F., Luo, Y. and Geyer, C.R. (2016) RecA inhibitors potentiate antibiotic activity and block evolution of antibiotic resistance. Cell Chem. Biol., 23, 381–391.

56. Nieto, J.M. and Juárez, A. (1999) The putative Orf4 of broad-host-range conjugative plasmid R446 could Be related to the H-NS family of bacterial nucleoid-associated proteins. Plasmid, 41, 125–127.

57. Francia, M.V., Varsaki, A., Garcillán-Barcia, M.P., Latorre, A., Drainas, C. and de la Cruz, F. (2004) A classification scheme for mobilization regions of bacterial plasmids. FEMS Microbiol. Rev., 28, 79–100.

58. Garcillán-Barcia, M.P., Francia, M.V. and de la Cruz, F. (2009) The diversity of conjugative relaxases and its application in plasmid classification. FEMS Microbiol. Rev., 33, 657–687.

59. Smillie, C., Garcillán-Barcia, M.P., Francia, M.V., Rocha, E.P.C. and de la Cruz, F. (2010) Mobility of plasmids. Microbiol. Mol. Biol. Rev., 74, 434–452.

60. Navarre, W.W., Porwollik, S., Wang, Y., McClelland, M., Rosen, H., Libby, S.J. and Fang, F.C. (2006) Selective silencing of foreign DNA with low GC content by the H-NS protein in Salmonella. Science, 313, 236–238.

61. Lang, B., Blot, N., Bouffartigues, E., Buckle, M., Geertz, M., Gualerzi, C.O., Mavathur, R., Muskhelishvili, G., Pon, C.L., Rimsky, S., et al. (2007) High-affinity DNA binding sites for H-NS provide a molecular basis for selective silencing within proteobacterial genomes. Nucleic Acids Res., 35, 6330–6337.

62. Bates, S., Roscoe, R.A., Althorpe, N.J., Brammar, W.J. and Wilkins, B.M. (1999) Expression of leading region genes on IncI1 plasmid ColIb-P9: genetic evidence for single-stranded DNA transcription. Microbiology, 145 (Pt 10), 2655–2662.

63. Gillings, M., Boucher, Y., Labbate, M., Holmes, A., Krishnan, S., Holley, M. and Stokes, H.W. (2008) The evolution of class 1 integrons and the rise of antibiotic resistance. J. Bacteriol., 190, 5095–5100.

64. Takeuchi, N., Hamada-Zhu, S. and Suzuki, H. (2023) Prophages and plasmids can display opposite trends in the types of accessory genes they carry. Proc. Biol. Sci., 290, 20231088.

65. Konno, N., Maeno, S., Tanizawa, Y., Arita, M., Endo, A. and Iwasaki, W. (2024) Evolutionary paths toward multi-level convergence of lactic acid bacteria in fructose-rich environments. Commun. Biol., 7, 902.

66. Rolff, J., Bonhoeffer, S., Kloft, C., Leistner, R., Regoes, R. and Hochberg, M.E. (2024) Forecasting antimicrobial resistance evolution. Trends Microbiol., 32, 736–745.

67. Ellabaan, M.M.H., Munck, C., Porse, A., Imamovic, L. and Sommer, M.O.A. (2021) Forecasting the dissemination of antibiotic resistance genes across bacterial genomes. Nat. Commun., 12, 2435.

68. Konno, N. and Iwasaki, W. (2023) Machine learning enables prediction of metabolic system evolution in bacteria. Sci. Adv., 9, eadc9130.

